# Molecular characterization of flower bud dormancy under constrained temperatures unveils a shallow dormancy stage induced by cold deprivation

**DOI:** 10.1101/2025.06.27.661981

**Authors:** Mathieu Fouché, Hélène Bonnet, Sylvain Prigent, Bénédicte Wenden

**Affiliations:** INRAE, Univ. Bordeaux, UMR Biologie du Fruit et Pathologie 1332, Villenave d’Ornon, France; Bordeaux Metabolome, MetaboHUB, PHENOME-EMPHASIS, 33140 Villenave d’Ornon, France

**Keywords:** Bud dormancy, Constrained temperature, Transcriptomics, Sweet cherry, Climate change, Shallow dormancy

## Abstract

Mild winters are becoming increasingly common in temperate regions due to climate change, which may have important impacts on ecosystems and agriculture. In particular, rising temperatures affect the progression of winter dormancy—a crucial developmental stage in perennial plants—making tree development and reproduction particularly vulnerable to climate change. A better understanding of how future temperature conditions will disrupt dormancy in cultivated fruit trees is crucial for anticipating the impeding consequences and identifying potential adaptation strategies. We investigated the effect of very constrained temperature conditions, i.e. several levels of cold deprivation and early cold exposure, on sweet cherry flower buds during dormancy onset and maintenance, using phenological observations and transcriptomic analyses.

We show that temperature is a major driver of dormancy progression as cold deprivation and early cold exposure strongly modify the timing of phenological phases as well as gene expression patterns. We identified genes and signaling pathways specifically activated and/or repressed by cold temperatures, and therefore potentially involved in the optimal progression of dormancy. Finally, thanks to an integrative analysis of molecular data obtained under natural and prolonged warm conditions, we characterize a distinct shallow dormancy phase induced by cold deprivation, with a unique molecular signature.

**Highlights:** Our phenological and molecular analysis of sweet cherry dormancy under constrained conditions reveals a gene expression timeline in response to temperature and uncovers a shallow dormancy stage induced by cold deprivation.

## Introduction

Many temperate perennial species undergo a wintering dormant period to protect bud tissues from unfavorable winter conditions for optimal vegetative bud break and blooming, which subsequently ensures survival and reproductive success (Coville, Frederick, 1920; Couvillon and Erez, 1985). Although a complex array of environmental conditions seems to regulate winter dormancy in trees, there is a general agreement that cold and mild temperatures are the main drivers of onset, maintenance and release of bud dormancy, especially in fruit trees (Arora *et al*., 2003; Horvath, 2009; Rohde *et al*., 2011; Chuine *et al*., 2016; Nilsson, 2022). This strong dependence to temperature conditions, and particularly chill requirements, prevents buds from precocious budbreak under false spring events, which are early warm conditions preceding a late cold spell (Gu *et al*., 2008; Chamberlain *et al*., 2019). Species that require a prolonged exposure to cold temperatures before responding to growth conditions (i.e. high chill requirements) will avoid leaf-out or flowering too early during periods of warm temperatures. However the downside is that the lack of chilling under mild winters results in abnormal patterns of budbreak, delayed flowering, bud necrosis and organ malformation (Bonhomme *et al*., 2005; El Yaacoubi *et al*., 2014; Funes *et al*., 2016). These impacts have already been observed in fruit trees (Erez, 2000; Jackson, 2000; Zguigal *et al*., 2006; Caffarra *et al*., 2011; Laube *et al*., 2014; Legave *et al*., 2015) but this trend is expected to worsen in the next decades due to global temperature increase (IPCC 6th report). Process-based modeling studies suggest that the lack of cold accumulation is responsible for local population extinction at the southern range margins of species (Morin, 2007). Furthermore, projections under future climatic scenarios expose high risks of severe cold deprivation for fruit and nut trees across growing regions (Luedeling and Brown, 2011; Campoy *et al*., 2019; Fernandez *et al*., 2023) where producers will need help to prepare for the impact of climate change.

Several studies have focused on the effects of mild winter conditions on fruit tree physiology and metabolism, usually submitting the trees to low, but not null, amounts of chill accumulation (Rakngan *et al*., 1996; Lecomte, I; Faye, F and Le Floc’h, F, 1998; Bonhomme *et al*., 2005; Yamane *et al*., 2008; Yamamoto *et al*., 2010). In particular, they found that severe cold deprivation led to necrosis in peach and pear flower buds and determined that bud necrosis could be linked to the inability to use carbohydrates and to abnormal functional water movement (Bonhomme *et al*., 2005; Yamamoto *et al*., 2010). However, these striking effects of cold deprivation on bud dormancy have barely been explored at the molecular level. Thanks to advances in sequencing technology in the last decade, transcriptomic and epigenetic analyses on fruit trees grown under natural environments, i.e. with enough exposure to cold temperatures, have led to the identification of key molecular regulators and signaling pathways associated with dormancy regulation (e.g. Ruttink *et al*., 2007; Bai *et al*., 2013; Vimont *et al*., 2019). In particular, these studies have highlighted a central role for *DORMANCY ASSOCIATED MADS-BOX* (*DAM*) genes in *Prunus* species, potentially through the regulation of phytohormones such as gibberellins (GA) and abscisic acid (ABA) during dormancy onset, maintenance and release (Bielenberg *et al*., 2008a; Zhu *et al*., 2015; Vimont *et al*., 2019; Yamane *et al*., 2019; Yu *et al*., 2020; Yang, 2021). . In particular, ABA was found to be implicated in dormancy onset and release through the regulation of transport capacity (Rinne *et al*., 2011; Tylewicz *et al*., 2018; Singh *et al*., 2019). In addition, oxidative stress and redox status could be a pivotal signal for dormancy release through the ROS production during endodormancy (Beauvieux *et al*., 2018) and previous transcriptomic studies conducted during bud dormancy in *Prunus* species have identified candidate genes related to oxidative stress such as oxidoreductase and peroxidases (Kuroda, 2005; Pérez *et al*., 2008; Yu *et al*., 2020). Moreover, carbohydrate metabolism has been found to change during dormancy progression, and the budburst capacity has been described to be link with its supply in carbohydrates and the activity of plasma membrane transporters after dormancy release (Beauvieux *et al*., 2018). In addition, sucrose which is considered to be both a storage and a protection against cold and desiccation increases in low temperature conditions due to the sucrose phosphate synthase activity (Cooke *et al*., 2012).

However, despite the well documented molecular actors involved in dormancy progression from all the past studies, little is known about the impact of increasing temperatures on biological pathways and the associated gene patterns throughout bud dormancy. One of the reasons for this knowledge gap is that these studies were rarely conducted in controlled conditions and therefore the response to extreme conditions expected under future climatic scenarios has not yet been investigated.

Exploring how signaling pathways and key genes change under constrained temperature conditions could help anticipating how global warming will affect bud dormancy and thus productivity for fruit trees. The main objective of this work was to investigate the effect of several levels of cold deprivation and precocious cold exposure on mature sweet cherry trees. Using phenological observations and transcriptomic analyses, this original study highlights how flower buds respond to temperature during dormancy, with major impacts on bud development and gene expression.

## Materials and methods

### Plant materials and experimental design

We used mature sweet cherry (*Prunus avium* L.) trees grown in 100L pots under field conditions in the INRAE campus (Villenave d’Ornon, France; 44.788996 N - 0.577947 W), with daily irrigation and nutrition. All experiments were conducted on the ‘Regina’ cultivar, which is characterized by high chill requirements and late flowering.

Two different experiments were set up to explore the response of temperature during sweet cherry dormancy. The first one was designed to establish several levels of cold deprivation during dormancy. The experiment started at the end of July 2018 after the flower initiation and before cooler nights in order to avoid chill accumulation. After bud set and flower initiation when flower buds are differentiated and begin their growth, four trees were placed in a walk-in climate chamber at 20°C during the day and 16°C during the night, under short day conditions (8h light/16h dark). Short day conditions were set in order to demonstrate the cold deprivation effect on dormancy under inducible dormancy photoperiod. Trees were then transferred from the climate chamber to natural conditions on October 31^th^ (CC2NOV), January 2^nd^ (CC2JAN), February 28^th^ (CC2MAR), one tree was left in the climate chamber until April 30^th^ (CC), after 93 days, 156 days, 213 days and 274 days under cold deprivation conditions, respectively (Fig. 1A). Two potted trees were left outside under natural environment as control.

**Figure 1.**
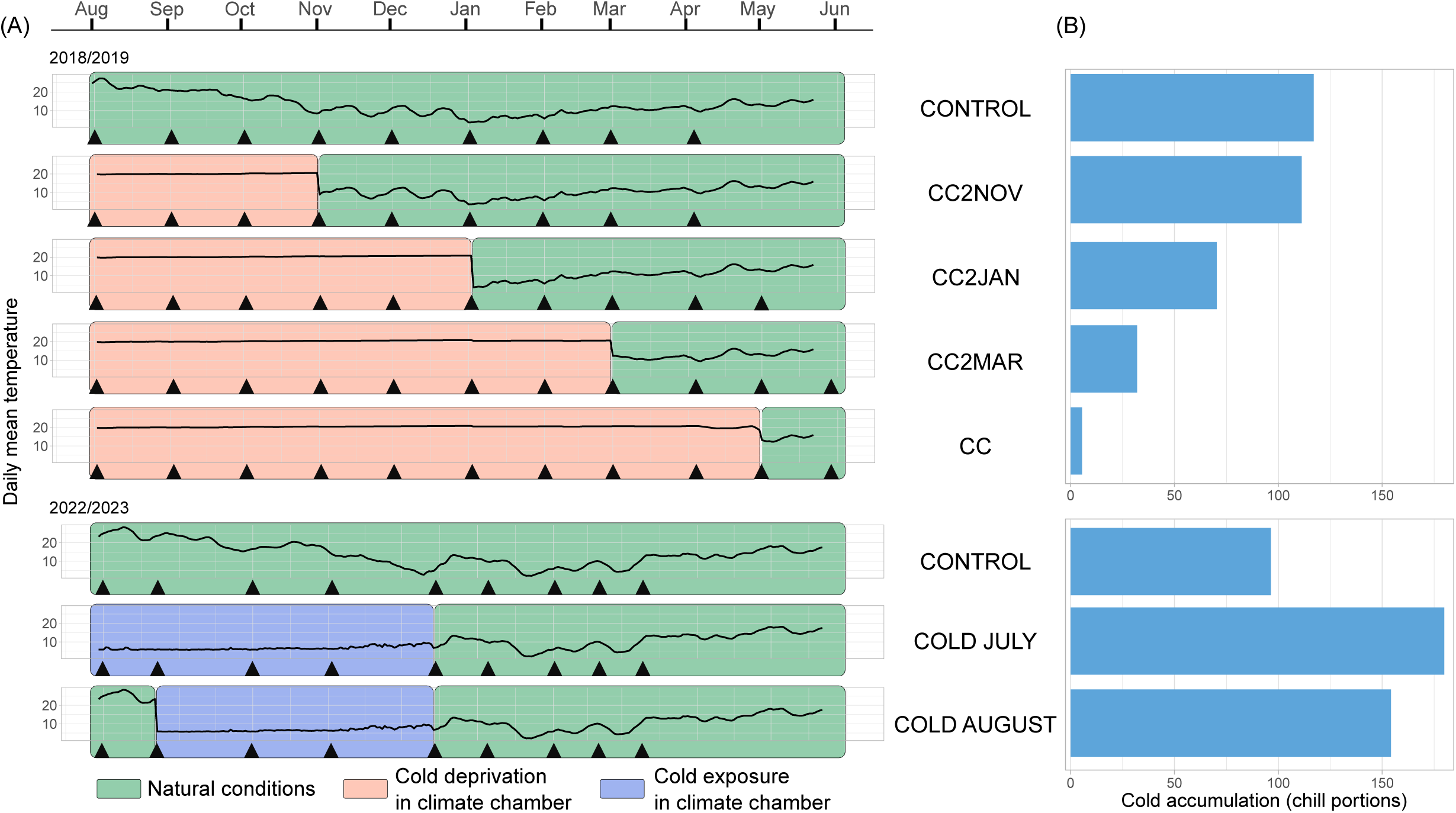
Experimental design for the study of potted trees under controlled temperature treatments. (A) Temperature conditions and sampling timepoints for trees submitted to different environmental conditions. The black lines represent the daily mean temperatures under each treatment. Green background indicates periods when potted trees were under field condition, red background indicates periods when potted trees were under cold deprivation in the climate chamber (20°C during the day, 16°C during the night, short days 8h light/16h dark). Trees were transferred from the climate chamber to natural conditions on October 31^th^ (CC2NOV), January 2^nd^ (CC2JAN), February 28^th^ (CC2MAR), one tree was left in the climate chamber until April 30^th^ (CC). Blue background indicates when potted trees were under cold exposure in the climate chamber (9°C during the day and 6°C during the night, long days 16h light/8h dark). Four trees were transferred to the climate on July 22th (COLD JULY) and four additional trees on August 21th (COLD AUGUST). Triangles indicate flower buds sampling timepoints for RNA-seq analysis for each treatment. (B) Total cold accumulated by the trees under each temperature treatment, calculated in chill portions using the dynamic model (Fishman et al., 1987) based on hourly temperatures recorded in the climate chamber and the orchard.

In contrast, a second experiment in controlled conditions was performed from the end of July 2022 to March 2023 to expose trees to early prolonged cold conditions, just after flower initiation. Four trees were transferred to the climate chamber at 9°C during the day and 6°C during the night under long days (16h light/8h dark) on July 22th (COLD JULY) and four additional trees on August 21th (COLD AUGUST) to create two cold exposure levels. Long day conditions were set in order to demonstrate the cold exposure effect on dormancy during non-inducible dormancy photoperiod. Two trees were left outside under natural environment as control.

Flower buds were sampled monthly until flowering date (Fig. 1A) in three biological replicates, corresponding to independent buds collected from one or several trees in each temperature treatment. In details, for all cold deprivation conditions (CC2NOV, CC2JAN, CC2MAR and CC), flower bud replicates were combined from all trees present in the climate chamber. Buds were flash frozen in liquid nitrogen and stored at − 80 °C prior to performing RNA extraction.

### Phenological observations

At the end of summer and in autumn, senescence was monitored and we considered senescence achieved when 50% of the leaves were either fallen or decolored (BBCH95, (Fadón *et al*., 2015). Spring phenology was monitored for budbreak (BBCH55), beginning of flowering (10% open flowers, BBCH61), full flowering (50-75% open flowers, BBCH65) and end of flowering (10% of flowers left on the branches, BBCH69). The flowering rate was used to evaluate the quality of flowering, i.e. the proportion of open flower related to the total number of flower buds on the branches.

### Temperature monitoring and calculation of chill accumulation

Hourly temperatures were recorded using Hobo pendant temperature/light 64k data logger (ref UA-002-64) and Omega data logger (ref OM-EL-USB-2).

Chill accumulation was estimated in chill portions (CP) using the dynamic model with previously published parameters (Fishman *et al*., 1987). The chill portions were summed from August 1st to June 30th.

### RNA extraction and paired-end (150pb) library preparation

Total RNA was extracted from approximately 100 mg of frozen powder obtained from grinded flower buds using RNeasy Plant Mini kit (Qiagen) with a minor modification: 1.5% PVP-40 was added in the extraction buffer RLT. RNA quality was evaluated using Tapestation 4200 (Agilent Genomics). Library preparation was performed on 1 μg of high-quality RNA (RNA integrity number equivalent superior or equivalent to 8.5) using the TruSeq Stranded mRNA Library Prep Kit High Throughput (Illumina cat. no. RS-122-2103). DNA quality from libraries was evaluated using fragment analyzer. The libraries were sequenced on a HiSeq3000 (Illumina), at the sequencing facility Get-Plage (Castanet-Tolosan, France).

### Mapping and differential expression analysis

The raw reads obtained from the sequencing were analysed using several publicly available softwares and in-house scripts. The quality of the reads was assessed using FastQC and possible adaptor contaminations and low quality trailing sequences were removed using Trimmomatic 0.36 (Bolger *et al*., 2014) with following settings fa:2:10:5:1 LEADING:3 TRAILING:3 SLIDINGWINDOW:4:15 MINLEN:36. Trimmed reads were mapped to the sweet cherry ‘Regina’ reference genome v.1 (Le Dantec *et al*., 2020) using STAR (Dobin *et al*., 2013). Raw read count was computed using HTSeq count (Anders *et al*., 2015) (Supplementary Table 1) and TPM (Transcripts Per Million) values were calculated with a house made R script inspired by (Li *et al*., 2010). We performed a differential expression analysis on raw read counts to identify expression patterns that changed during dormancy. First, data were filtered by removing lowly expressed genes (average read count < 3) and genes not expressed in most samples (read counts = 0 in more than 75% of the samples). Then, differentially expressed genes (DEGs) between either each time point or cold deprivation conditions were assessed using DEseq2 R Bioconductor package (Love *et al*., 2014), in the statistical software R (R Core Team 2022). Genes with an adjusted p-value (padj) < 0.05, using the Benjamini-Hochberg multiple testing correction method, and a log2 fold change < -1 or > 1 between at least two dates or conditions were assigned as DEGs.

### Principal component analysis and hierarchical clustering

Principal component analyses (PCA) were performed on TPM values of the samples using the R packages FactoMineR and factoextra. The *HCPC* function was used to create sample’s dendrogram from the PCA results. Distances between the samples were calculated based on Pearson’s correlation on TPM values. We applied a hierarchical clustering analysis using the *hclust* function on the distance matrix to define gene expression clusters.

### Gene ontology enrichment analysis

The sweet cherry ‘Regina’ reference genome was annotated using Blast2GO (Gotz *et al*., 2008) for gene ontology (GO) terms. Using the topGO package for R (Alexa A, Rahnenfuhrer J, 2023), we performed an enrichment analysis on GO terms for Biological Processes and for Molecular Function and Cellular Component on the DEGs compared to the whole set of annotated genes, and for each cluster, compared to the total DEG list, based on a classic Fisher algorithm. Enriched GO terms were filtered with a p-value < 0.05 and the ten GO terms with the lowest p-values were selected for representation.

### Predictive transcriptomic

Glmnet package (Friedman *et al*., 2010) that fits generalized linear and similar models via penalized maximum likelihood was used to perform dormancy stage prediction based on gene expression level.

The Glmnet algorithms use cyclical coordinate descent, which successively optimizes the objective function over each parameter with others fixed, and cycles repeatedly until convergence. For each dormancy stage, 100 simulations were run with different combinations of training (80% of data) and testing (20% of data) datasets created using stratified sampling based on the phenotype to predict. Genes were considered as marker genes for one dormancy stage when it was present in more than 75% of the models. During the training phase, a grid of 40x40 was used to determine the best hyper parameters using a 3 times 10-fold cross-validation. The best models were chosen based on maximization of the accuracy during the training step.

## Results

### Constrained temperatures alter the timing of phenological phases

Four levels of cold deprivation under short days, corresponding to a range from 111 to 5.5 CP accumulated (Fig. 1B), led to a range of delays in leaf fall and flowering dates (Fig. 2A). The control trees grown under natural conditions showed a beginning of senescence at the end of October, and were leafless on November 30^th^ (Fig. 2A). In comparison, under short days and cold deprivation treatments, senescence occurred two months later around mid-January. Moreover, trees treated with cold deprivation flowered at a later date than the control trees, with an increasing delay associated with a longer cold deprivation treatment (Fig. 2A). Indeed, when the control trees and the tree transferred from the climate chamber to field conditions in November both showed flowering dates around March 27^th^ (Fig. 2A), flowering dates were markedly delayed in trees that were not subjected to cold conditions. For example, the potted tree submitted to cold deprivation until March (CC2MAR), and which had accumulated around 32 CP, flowered on June 24^th^, corresponding to three months delay compared to the trees grown under control condition (Fig. 2A). In contrast, trees submitted to an early exposure to cold in July and August showed early senescence, occurring 51 days and 61 days after transfer to cold, respectively. Flowering was also impacted for these trees that flowered 29 days and 22 days earlier than the control, respectively (Fig. 2A).

**Figure 2.**
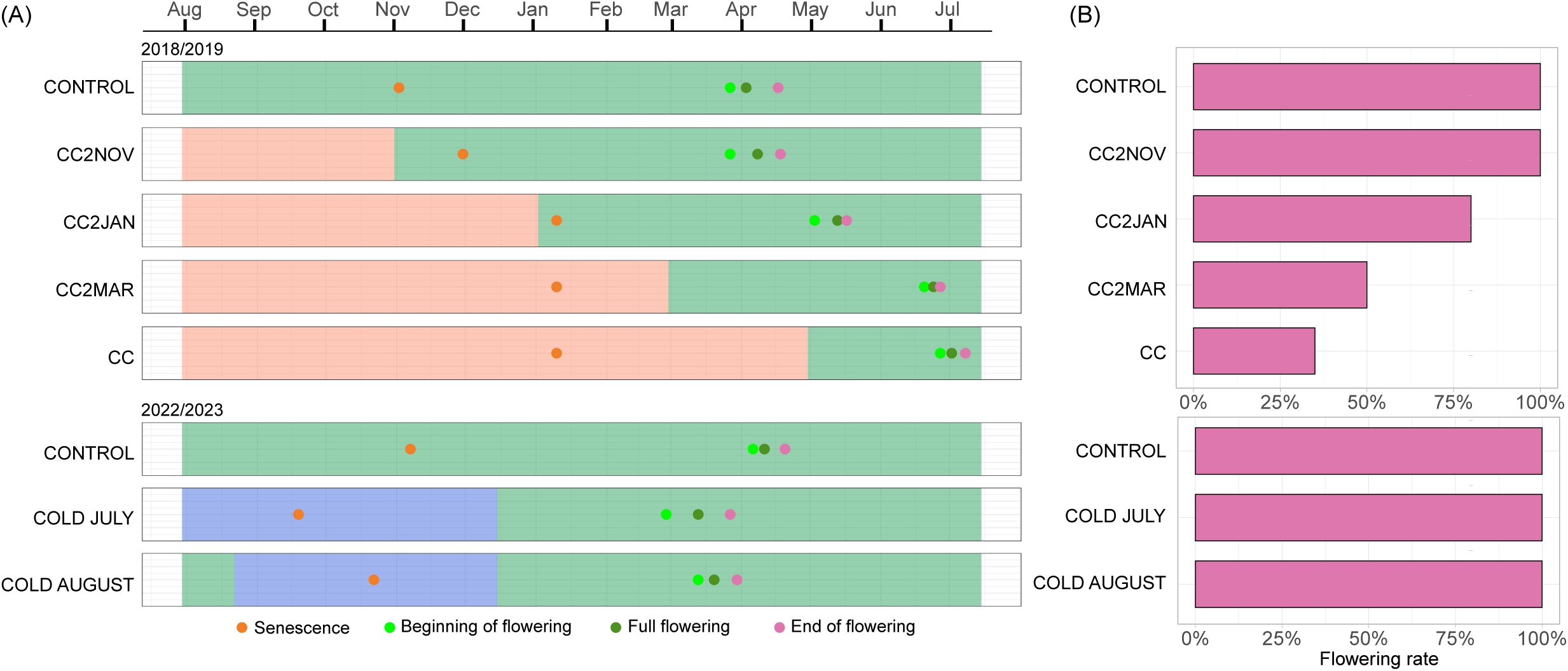
Phenological changes induced by constrained temperature conditions. (A) Senescence and flowering dates recorded for trees submitted to each treatment. Green background indicates periods when potted trees were under field condition, red background indicates periods when potted trees were under cold deprivation in the climate chamber (20°C during the day, 16°C during the night, short days 8h light/16h dark). Trees were transferred from the climate chamber to natural conditions on October 31^th^ (CC2NOV), January 2^nd^ (CC2JAN), February 28^th^ (CC2MAR), one tree was left in the climate chamber until April 30^th^ (CC). Blue background indicates when potted trees were under cold exposure in the climate chamber (9°C during the day and 6°C during the night, long days 16h light/8h dark). Four trees were transferred to the climate on July 22th (COLD JULY) and four additional trees on August 21th (COLD AUGUST) (B) Flowering rate observed for trees under the different temperature treatments.

It has been observed that flowering quality, i.e. the percentage of open flowers and morphological abnormalities, was reduced in fruit trees under warm winters (Tominaga *et al*., 2022). We investigated to which extend flowering quality was impacted under severe cold deprivation. We found that the percentage of open flowers decreased with the length of the cold deprivation treatment, from 100% for CC2NOV to 35 % for the most severe cold deprivation condition (CC), while early cold exposure did not affect the flowering rate and quality (Fig. 2B). Moreover, flowers that have been observed after cold deprivation showed abnormal formation such as deformed verticilles, with potential impact on the reproductive efficiency (Supplementary Fig. 1C).

### Bud dormancy stages are characterized by specific gene expression patterns under natural conditions

In order to understand how and when molecular mechanisms are involved in dormancy progression under different temperature treatments, we first explored gene expression levels over the dormancy period under normal environmental conditions as a baseline for further analyses. This was achieved based on a transcriptomic analysis by RNA-seq conducted on flower buds sampled from the control trees. In order to characterize the genes and signaling pathways specifically activated during the different stages of dormancy on field conditions, we performed a PCA on the TPM values for the 10,130 identified DEGs (Supplementary Table 2). When projected on the first two PCA dimensions, we found that the first two axes PCA1 and PCA2 allowed discriminating the biological replicates by dates into three main groups, defined using hierarchical clustering (Fig. 3B). Based on the sampling dates and the definition of dormancy stages proposed by Lang et al. (1987), we associated these groups with paradormancy, endodormancy and ecodormancy (Fig. 3A). First, samples from July to October were classified as paradormancy, which corresponds to the early flower bud stages, including flower primordia organogenesis in the summer, followed by reduced growth and dormancy onset via inhibited correlations from active organs in combination with cooler temperature at the beginning of fall. Samples from November to February correspond to endodormancy, when growth arrest is controlled by endogenous inhibitors to protect vital tissues from frost and buds cannot develop even under growth conditions. A prolonged exposure to cold temperatures is needed to release the growth inhibition and break the endodormancy phase. Finally, the last cluster corresponds to February and March sampling dates, which is commonly defined as ecodormancy when buds are able to respond to warmer growth conditions. In order to explore the distinct gene expression patterns associated with the three dormancy stages, we performed a hierarchical clustering on DEGs for samples obtained under natural conditions. We identified six main clusters of gene expression patterns (Fig.3B), that we grouped into i) genes with a maximum expression level during paradormancy (cluster 3: 2589 genes and cluster 5: 1188 genes), ii) clusters with a maximum expression level during endodormancy (cluster 6: 1535 genes and cluster 2: 961 genes) and iii) clusters with a maximum expression level during ecodormancy (cluster 1: 1948 genes and cluster 4: 1909 genes).

**Figure 3.**
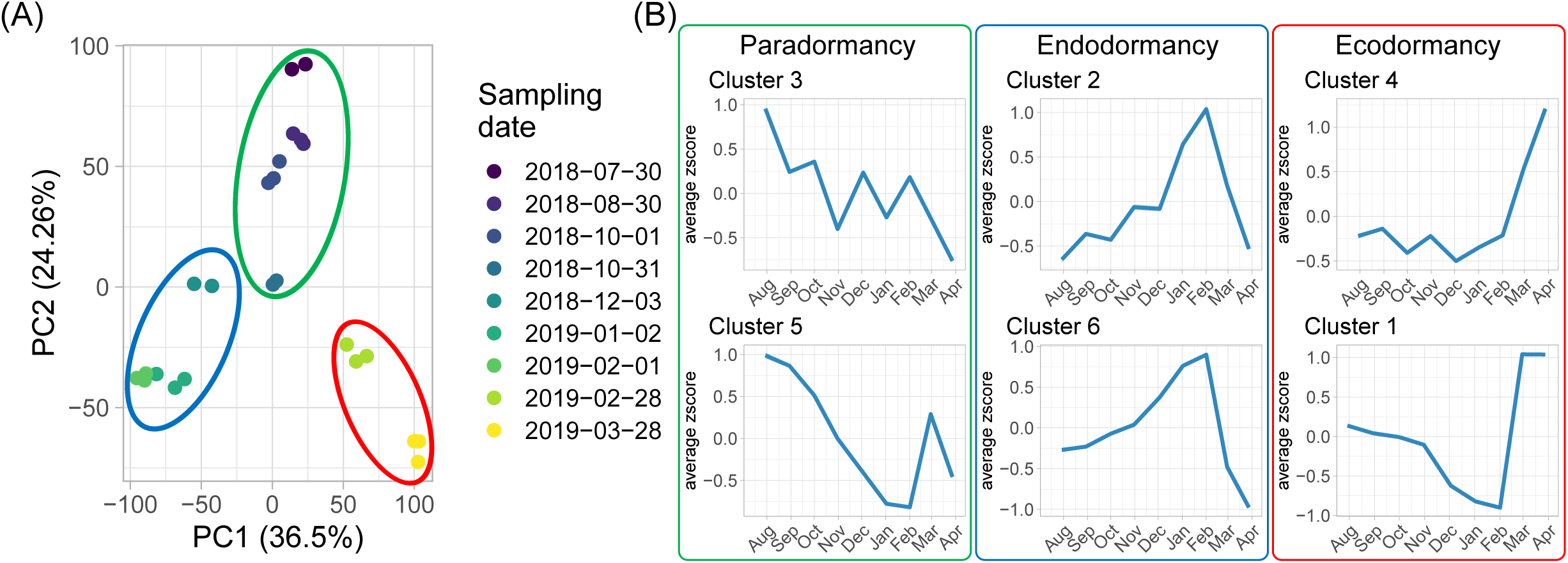
Gene expression levels accurately capture flower bud dormancy stages under natural conditions. (A) The principal component analysis was conducted on the TPM (transcript per millions reads) values for genes differentially expressed between sampling dates for flower buds sampled from sweet cherry ‘Regina’ potted trees grown under natural conditions. Samples in the green circle correspond to the paradormancy stage, samples in the blue circle correspond to the endodormancy stage, samples in the red circle correspond to the ecodormancy stage. Each point represents one biological replicate (n = 3 for each sampling date), corresponding to independent buds collected from two trees. (B) Average expression profile for genes belonging to the six clusters, organized according to the timing their expression peak in relation to the three main dormancy phases. Expression values are normalized from TPM values and *z-scores* are represented here.

### Genes highly expressed during paradormancy are characterized by two distinct responses to cold deprivation

Among the DEGs, 25% belonged to the cluster 3 corresponding to genes that were highly expressed in August during paradormancy and then down-regulated from September and endodormancy towards minimum levels during ecodormancy (Fig. 3B). Interestingly, under cold deprivation, the expression patterns for these genes remained high, with a slight increase throughout the treatment, and they were characterized by a sharp decrease after transfer to field conditions, with colder temperatures, showing that these genes were rapidly down regulated by the cold signal (Fig. 4A). This result was confirmed under cold exposure where genes belonging to cluster 3 were markedly repressed in response to the early cold signal while high expression was maintained under warmer natural conditions (July to November; Fig. 4A). We found that genes characterized by this specific inhibition by cold were significantly associated with RNA and mRNA metabolic process, mRNA splicing, and related to spliceosome (Fig. 4B). Interestingly, in addition to these pathways, we identified transcription factors related to dormancy regulation such as *DAM1* and *DAM6*, which are known to be expressed during paradormancy and downregulated during endodormancy and ecodormancy under natural conditions (Bielenberg *et al*., 2008b; Yamane *et al*., 2011; Vimont *et al*., 2019). We therefore confirmed that cold deprivation conditions maintained a high expression for *DAM1* and *DAM6* while they were promptly repressed under cold conditions upon transfer to field conditions or early cold exposure (Fig. 4C).

**Figure 4.**
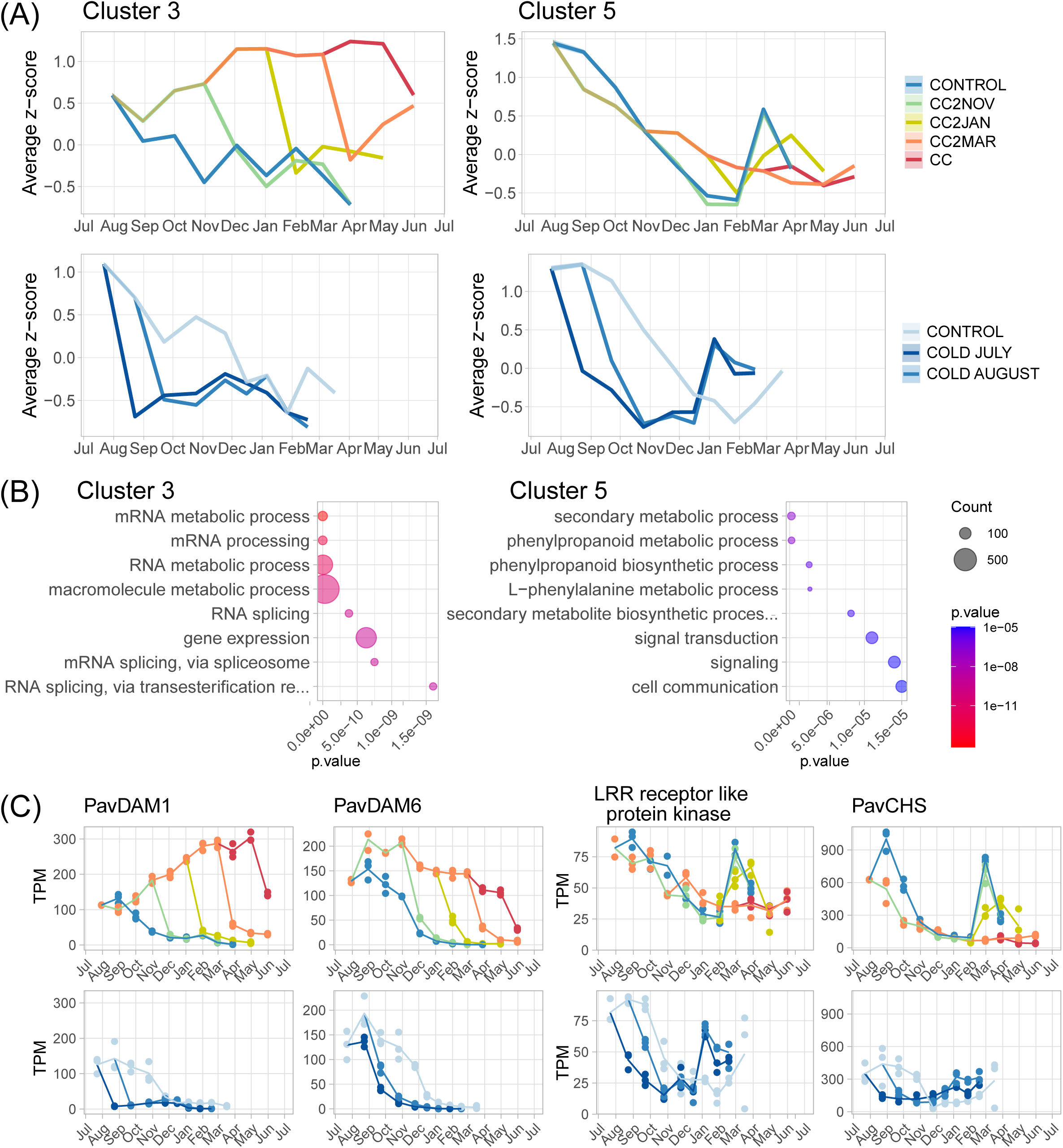
Molecular characterization of gene clusters associated with paradormancy. (A) Average expression profile for genes belonging to clusters 3 and 5 under cold deprivation and cold exposure treatments. Expression values are normalized from TPM values and *z-scores* are represented here. (B) Enrichment in Gene Ontology terms for Biological Processes for genes belonging to clusters 3 and 5. (C) Expression levels in TPM (transcripts per million reads) of genes of interest belonging to clusters 3 and 5 under cold deprivation and cold exposure treatments. Blue, green, yellow, orange and red lines correspond to Control, to CC2NOV, to CC2JAN, to CC2MAR and CC conditions for cold deprivation treatments, respectively and light blue line, medium blue line and blue line correspond to Control, to Cold August and to Cold July for cold exposure treatments, respectively. Dot size represents the number of genes belonging to the clusters associated with the GO term.

Unexpectedly, we observed that genes belonging to cluster 5, which are highly expressed during early paradormancy and gradually repressed towards endodormancy, were also down-regulated under cold deprivation conditions, albeit in a more moderate way (Fig. 4A). Although this remarkable inhibition of expression without cold signal could indicate that temperature is not the main regulating signal for genes belonging to cluster 5, their expression was rapidly repressed under early cold exposure (Fig. 4A), thus suggesting that both cold temperatures and endogenous signals might interact to control their expression. Furthermore, results indicated that a prolonged exposure to cold was necessary to induce a resumption of expression for genes belonging to cluster 5, as observed in February/March under natural conditions or earlier in December/January for trees exposed to an early cold signal (Fig. 4A), which may be associated with dormancy release and growth potential in ecodormancy. Our GO enrichment analysis revealed that these genes repressed then activated by prolonged cold were playing a role in signal transduction, cell communication and secondary metabolism such as phenylpropanoid (Fig. 4B). In this cluster 5, candidate genes were identified such as a *CHALCONE SYNTHASE* (*PavCHS*) gene, which plays a pivotal role in secondary metabolism or genes involved in signal transduction like *PavLRR receptor like serine/threonine-protein kinase* which belongs to a large gene family implicated in developmental processes via in particular hormone perception (Torii, 2004).

### Genes linked to endodormancy are sequentially activated and inhibited upon cold exposure

Genes highly expressed during endodormancy have been a focus of interest since they reveal the underlying processes occurring during an apparently inactive state. Consequently, we explore how these genes were expressed under very constrained temperature regimes. Genes belonging to the clusters 2 and 6 were highly expressed during endodormancy, potentially activated by decreasing temperatures at early endodormancy, and then inhibited following a certain cold accumulation threshold under control conditions (Fig. 3B). However, their responses under cold deprivation differed. Indeed, under cold deprivation, genes belonging to cluster 2 were maintained at a low level of expression while expression was high and stable for genes belonging to cluster 6. In addition, they were all rapidly up-regulated after transfer to natural conditions or cold exposure then down-regulated following a prolonged cold accumulation (Fig. 5A).

**Figure 5.**
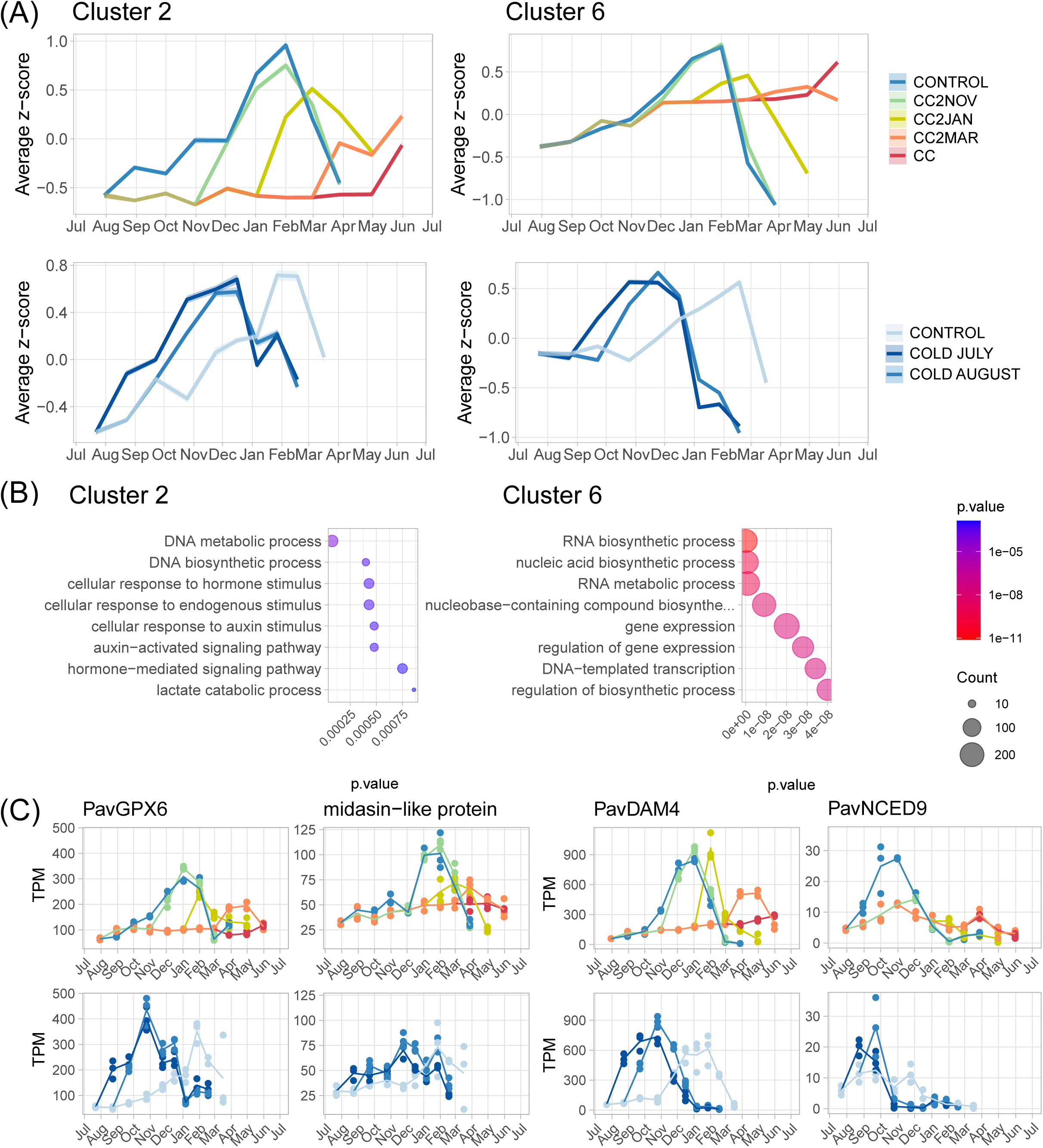
Molecular characterization of gene clusters associated with endodormancy. (A) Average expression profile for genes belonging to clusters 2 and 6 under cold deprivation and cold exposure treatments. Expression values are normalized from TPM values and *z-scores* are represented here. (B) Enrichment in Gene Ontology terms for Biological Processes for genes belonging to clusters 2 and 6. (C) Expression levels in TPM (transcripts per million reads) of genes of interest belonging to clusters 3 and 5 under cold deprivation and cold exposure treatments. Blue, green, yellow, orange and red lines correspond to Control, to CC2NOV, to CC2JAN, to CC2MAR and CC conditions for cold deprivation treatments, respectively and light blue line, medium blue line and blue line correspond to Control, to Cold August and to Cold July for cold exposure treatments, respectively. Dot size represents the number of genes belonging to the clusters associated with the GO term.

Our analysis of cluster 2 showed a specific enrichment in genes involved in DNA and regulation of cellular metabolic processes (Fig. 5B). Interestingly we also found that GO terms associated with glutathione transferase activity (Supplementary Table 3) were over-represented and we highlighted a gene coding for a glutathione peroxidase (*PavGPX6*) which was up-regulated during endodormancy and could participate in the antioxidant regulation in response to stress such as cold (Siller-Cepeda, 1992; Beauvieux *et al*., 2018). In this cluster, we also identified a transcription factor, which is a an homologue of *Atmidasin-like protein* that has been described to participate in ribosome biogenesis regulating plant growth and reproduction (Li *et al*., 2022) (Fig. 6C). This result is in line with the Cellular Component transcription factor elongation term found in the GO enrichment analysis (supplementary table 3). For cluster 6, our GO enrichment analysis revealed enrichment in genes implicated in the regulation of RNA biosynthetic process, gene expression, nucleic acid metabolic process (Fig. 5B). Accordingly, we found that the transcription factor *PavDAM4*, which has been previously described as a key player for dormancy maintenance (Zhu *et al*., 2020), was inhibited under cold deprivation conditions but sharply activated then inhibited upon exposure to cold conditions (Fig. 5C). Furthermore, we found that genes involved in ABA pathway were enriched in cluster 6 (Supplementary Table 3), such as *PavNCED9*, a key regulator of ABA biosynthesis (Fig. 5C). These results confirmed the key role of transcription factors such as *PavDAM4*, redox and ABA signalling in the regulation of endodormancy and we also found that DNA, RNA and protein metabolic processes were essential during endodormancy. These key signalling pathways are activated then inhibited under prolonged cold conditions.

**Figure 6.**
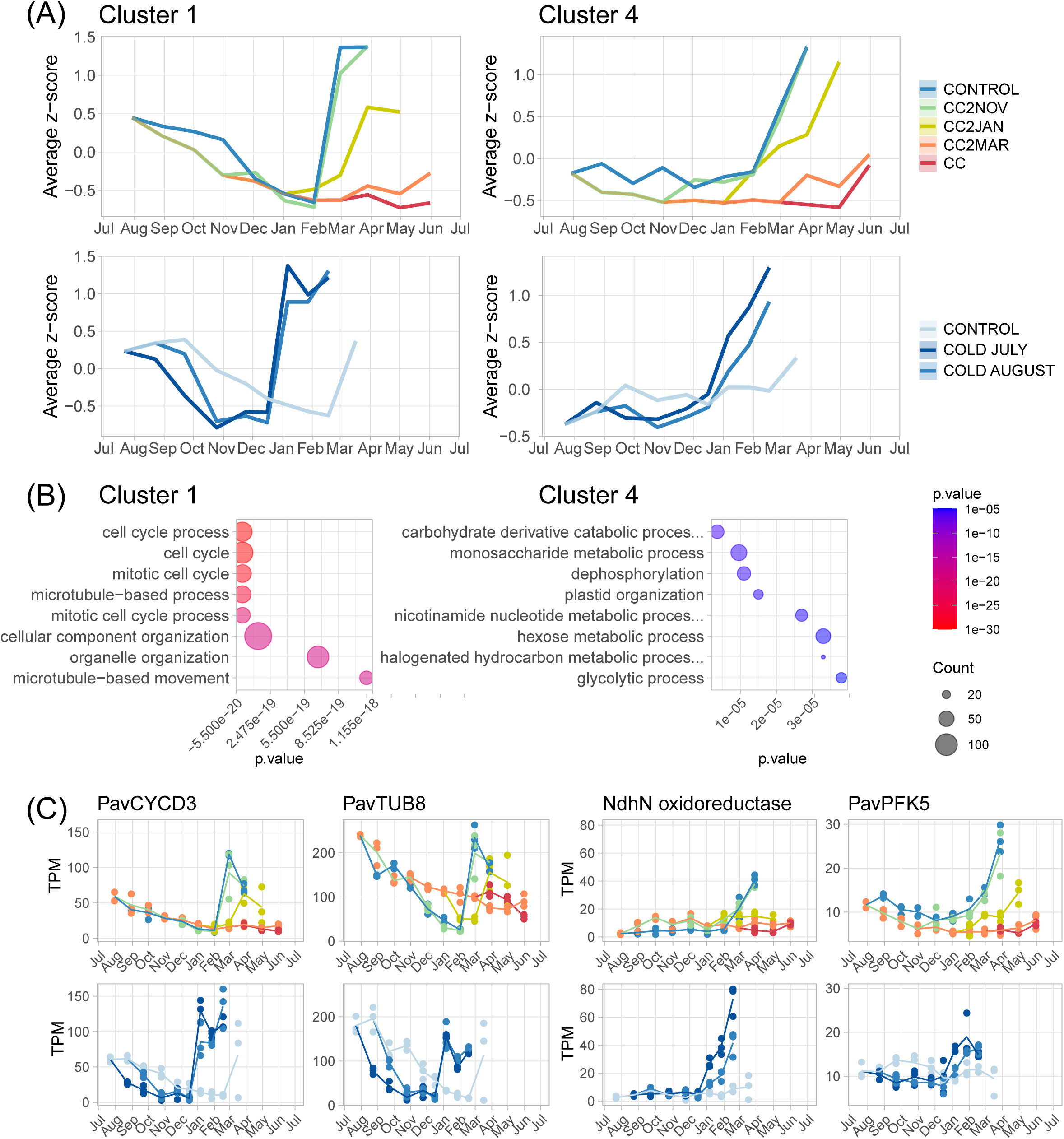
Molecular characterization of gene clusters associated with ecodormancy. (A) Average expression profile for genes belonging to clusters 1 and 4 under cold deprivation and cold exposure treatments. Expression values are normalized from TPM values and *z-scores* are represented here. (B) Enrichment in Gene Ontology terms for Biological Processes for genes belonging to clusters 3 and 5. (C) Expression levels in TPM (transcripts per million reads) of genes of interest belonging to cluster 1 and 4 under cold deprivation and cold exposure treatments. Blue, green, yellow, orange and red lines correspond to Control, to CC2NOV, to CC2JAN, to CC2MAR and CC conditions for cold deprivation treatments, respectively and light blue line, medium blue line and blue line correspond to Control, to Cold August and to Cold July for cold exposure treatments, respectively. Dot size represents the number of genes belonging to the clusters associated with the GO term.

### Genes highly expressed during ecodormancy may be activated by extended cold accumulation

Genes expressed during ecodormancy, as shown in clusters 1 and 4, are typically activated after several month of prolonged cold exposure (Fig. 6A), as confirmed by the patterns observed after an early transfer to cold conditions (COLD JULY and COLD AUGUST), where expression for ecodormancy genes increased in December and January, i.e. after five to six months of cold conditions.

In addition to a sharp expression increase toward the ecodormancy stage, genes belonging to cluster 1 were also expressed during early paradormancy then repressed during endodormancy. Interestingly, we found that the down-regulation, associated with endodormancy, occurred earlier for trees transferred to cold in July than for trees transferred in August, thus suggesting a cold-mediated inhibition of expression (Fig. 6A), similarly to the pattern observed for genes identified as paradormancy genes in cluster 5 (Fig. 4A).

We explored the signaling pathways activated during ecodormancy and we found enrichment in GO terms associated with cell cycle regulation, including DNA replication and cytoskeletal but also plastid organization and metabolic processes such as organonitrogen compound biosynthesis and glycolytic process (Fig. 6B). Likewise, we identified candidate genes in clusters 1 and 4 previously described to be potentially involved in the transition toward growth resumption (Fig. 6C). For instance in cluster 1, we found *PavCYCD3* which belongs to the cyclin gene family, described to regulate the transition between cell cycle phases (Menges *et al*., 2006). *PavTUB8*, also identified as a gene of interest, is homologous to genes involved in the microtubule formation, which participates in cell division regulated by microtubule cytoskeleton and driving the segregation of chromosomes and the formation of two daughter cells (Motta and Schnittger, 2021). The candidate gene approach in cluster 4 pinpointed an oxidoreductase gene that could contribute to the regulation of ROS associated with dormancy release (Beauvieux *et al*., 2018) and *PavPFK5*, homologous to a phosphofructose kinase which has been reported to be implicated in carbohydrate metabolism and plant adaptability to environmental changes (Fig. 6C). Overall, we found that genes activated during ecodormancy, potentially in response to a prolonged exposure to cold, were associated with growth and cell activity and their low expression may suggest that cell division and metabolic processes are very limited under extended cold deprivation conditions.

### A distinct dormancy stage is revealed under strict cold deprivation conditions

The analysis of expression patterns under different temperature regimes showed that cold deprivation conditions strongly constrained the expression levels of genes involved in dormancy onset, maintenance and release. In this context, we expected the trees to maintain a paradormancy stage in the absence of cold signal. However, trees under cold deprivation achieved senescence, although delayed compared to control conditions (Fig. 2A), and buds appeared dormant (Supplementary Fig. 1A). Therefore we further investigated the global molecular state of the samples based on the transcriptome. For this, we performed a PCA on DEGs expression levels from the samples obtained under cold deprivation and cold exposure conditions and we compared the projected coordinates between dates and conditions. While samples obtained under control conditions and cold exposure were well separated into the three dormancy stages (Fig. 7A-B), we found that samples from buds submitted to prolonged cold deprivation treatments were projected in close clusters, clearly differentiated from other dormancy stages, especially for the prolonged cold deprivation conditions (CC, December to April). Surprisingly, a further clustering analysis classed these samples into a separate cluster, related but distinct from paradormancy samples (Fig. 7C). These results highlight a specific dormancy stage, which could correspond to a prolonged paradormancy that we defined as shallow dormancy, in opposition to endodormancy (i.e. deep dormancy). From a phenological point of view, shallow dormancy is characterized by a very late senescence and bud set which occurred in the absence of cold. This result indicates that although cold temperatures are necessary to enter endodormancy, as observed in the control and CC2NOV conditions, other signals, such as endogenous cues, seem to be involved in the establishment of shallow dormancy. Interestingly, after transfer to cooler temperatures under natural conditions in May following nine months of cold deprivation, buds did entered an endodormancy stage, at least at the molecular level (CC_5; Fig. 7A,C). This result suggests that shallow dormancy is not irreversible since buds still responded to the cold signal.

**Figure 7.**
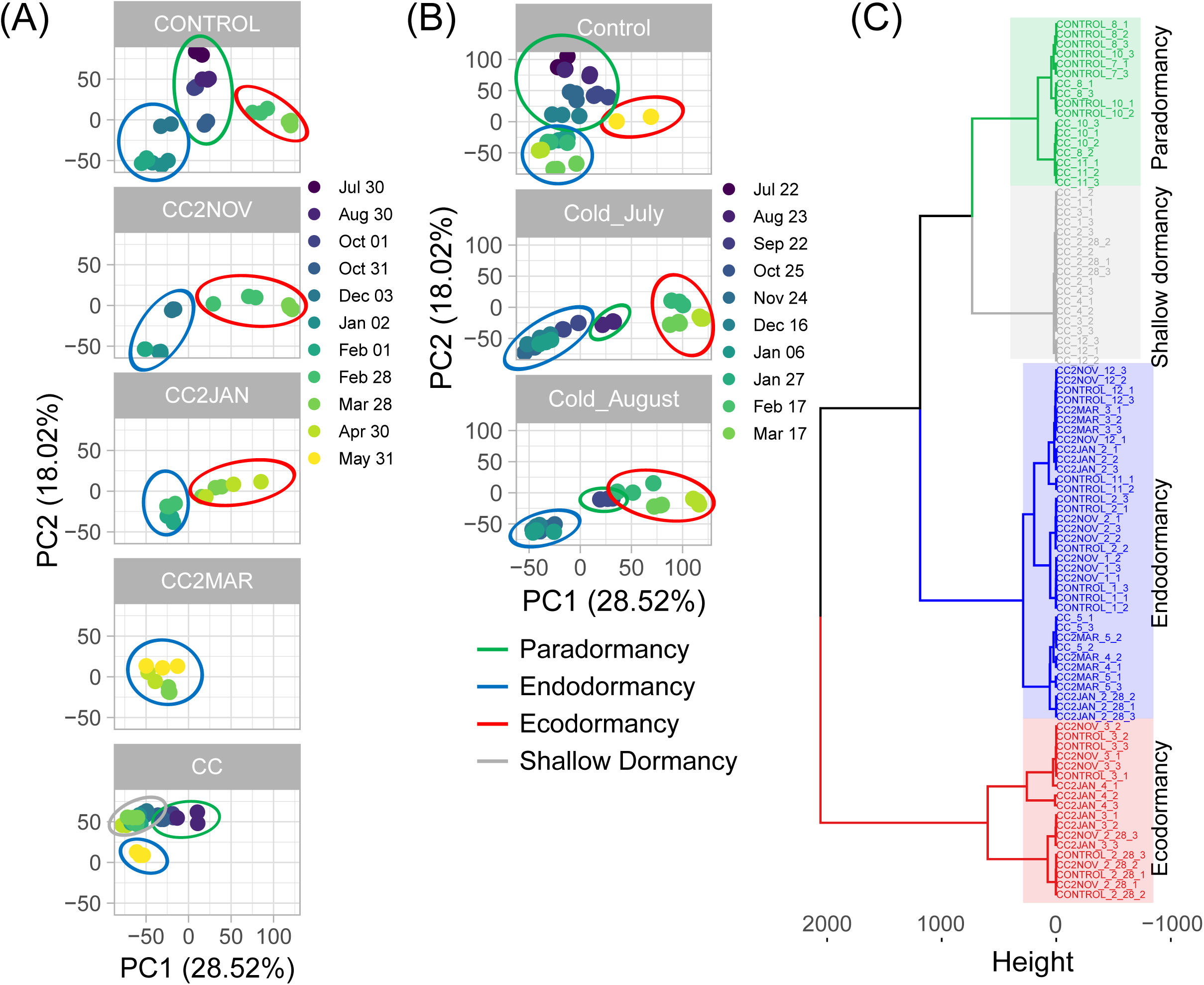
Gene expression levels obtained under constrained conditions unravel a singular bud dormancy stage. The principal component analysis was conducted on the TPM (transcript per millions reads) values for genes differentially expressed between sampling dates for flower buds sampled from sweet cherry ‘Regina’ potted trees grown under (A) cold deprivation and (B) cold exposure treatments. Samples in the green circle correspond to the paradormancy stage, samples in the blue circle correspond to the endodormancy stage, samples in the red circle correspond to the ecodormancy stage, samples in the grey circle belong to a fourth sample group, hereby defined as shallow dormancy stage based on a (C) hierarchical clustering on principal component analysis conducted on the cold deprivation experiment. Each point represents one biological replicate (n = 3 for each sampling date), corresponding to independent buds collected from one or several trees.

At the molecular level, based on a predictive transcriptomic model, we were able to pinpoint a reduced group of 116 marker genes that could significantly predict the shallow dormancy stage distinctly from other dormancy stages with an accuracy of 0.9988 (Fig. 8; Supplementary Table 4 ;Supplementary Table 5), thus validating its definition as a distinct bud stage. The majority of these marker genes belonged to the Cluster 3 (Fig. 8), previously described in Fig. 4 to be related to paradormancy stage and characterized by high expression levels under cold deprivation treatment.

**Figure 8.**
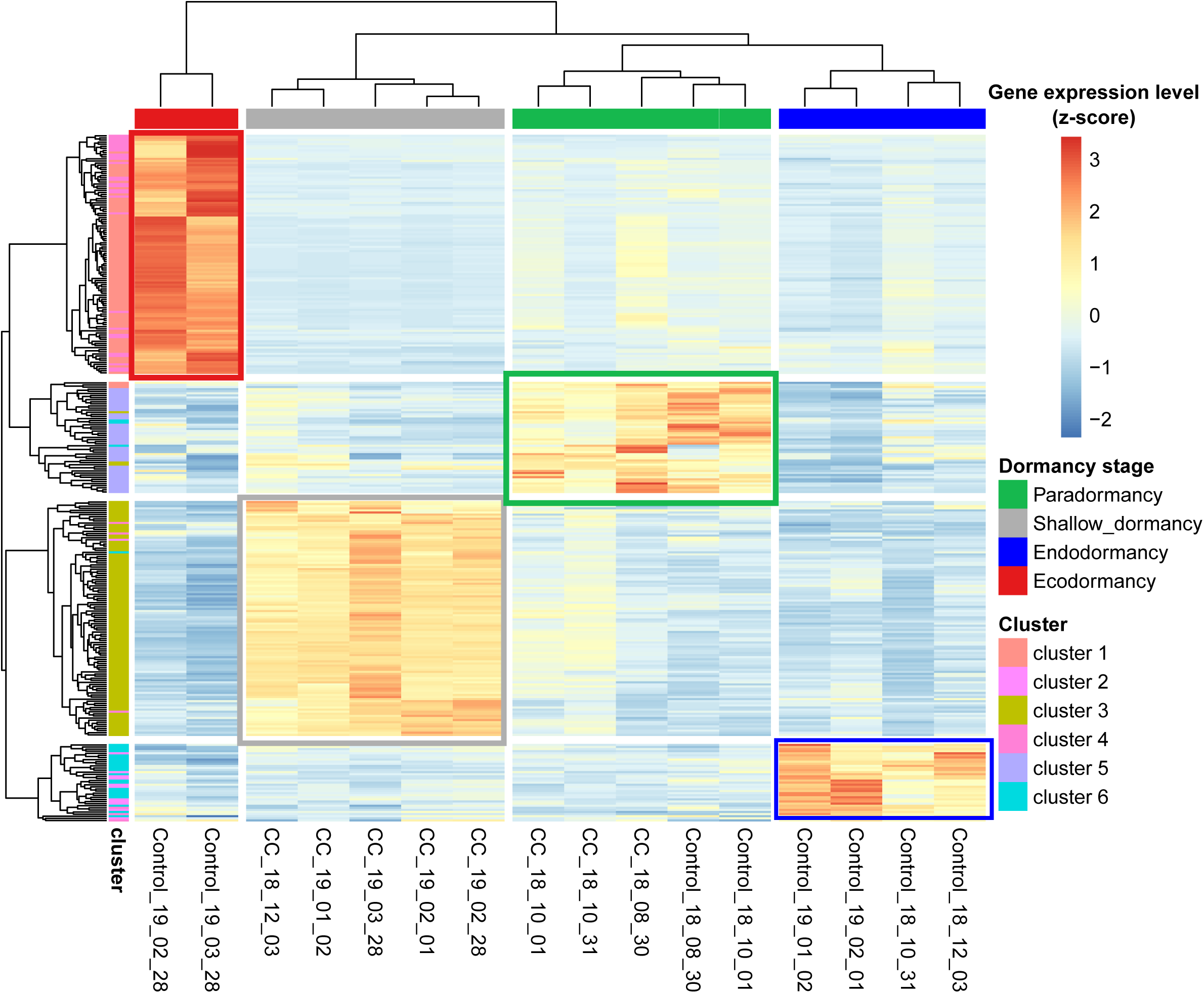
Dormancy stage associated with gene expression clusters. Heatmap indicating hierarchical clustering of genes identified by predictive transcriptomic (with at least an occurrence >75). Each column corresponds to the average gene expression for flower buds at a given date. Each row corresponds to the expression pattern across samples for one gene. Dormancy stage of each date is reported at the top of each column and cluster identification is indicated on the left of each row.

### Molecular characterization of the shallow dormancy stage induced by prolonged warm temperatures

Since we showed we could pinpoint shallow dormancy as a distinct stage based on whole transcriptome data (Fig. 7C), we investigated the signaling pathways that best characterize establishment and maintenance of this particular dormancy stage. For this, we focused only on data obtained in cold deprivation and we explored the specific expression patterns under these conditions, using a hierarchical clustering on the expression levels of DEGs for buds sampled from August to April in controlled conditions (CC). This analysis revealed two main expression patterns: genes that are down-regulated (2682 genes) or up-regulated (2664 genes) under prolonged cold deprivation (Fig. 9A; Supplementary Table 6).

**Figure 9.**
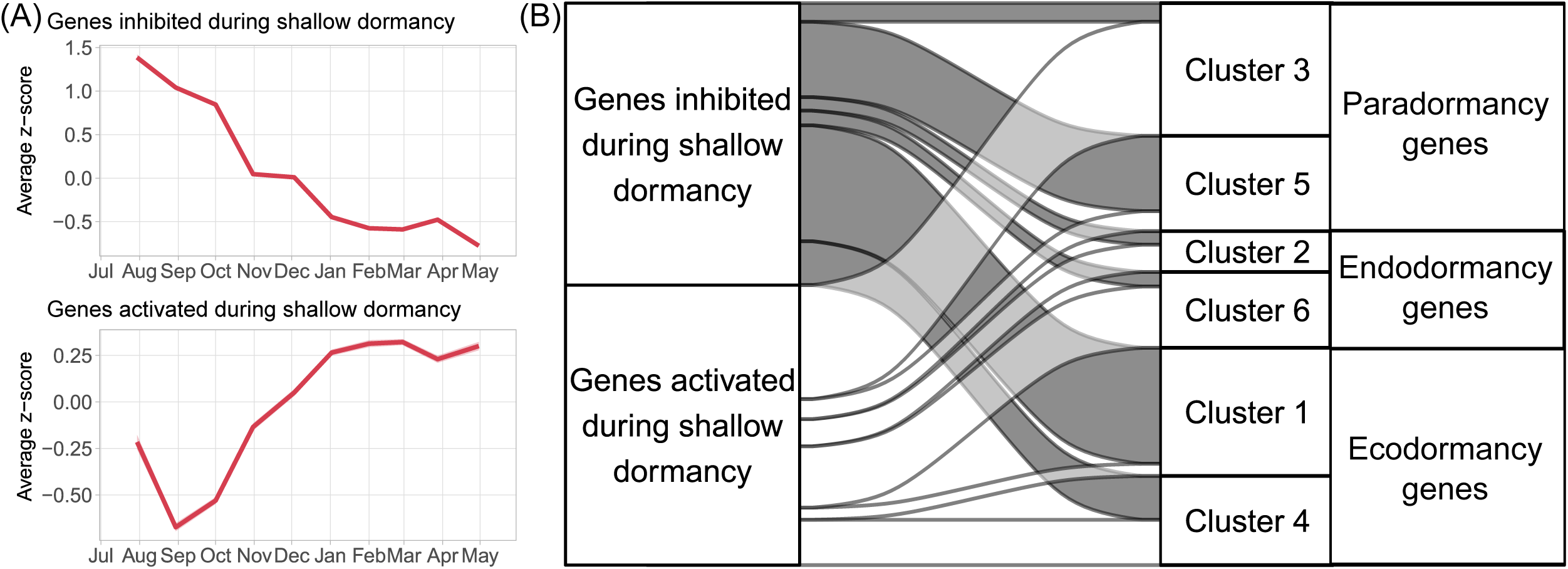
Molecular characterization of shallow dormancy. (A) Average expression profile for genes specifically activated and inhibited during severe cold deprivation. Expression values are normalized from TPM values and *z-scores* are represented here. (B) Cluster classification of genes inhibited or activated during shallow dormancy, based on clusters defined using natural conditions expression patterns.

We investigated how these genes were globally expressed under natural conditions to better understand which specific mechanisms were activated or repressed to induce and maintain shallow dormancy. We found that genes inhibited during shallow dormancy mostly belonged to clusters 5 and 1, associated with paradormancy and ecodormancy, respectively (Fig. 9B), and involved in cell cycle, cytoskeleton, organelle organization and secondary metabolic processes (Fig. 4B, Fig. 6B), which is in line with a gradual cessation of growth and cellular activity throughout shallow dormancy. On the opposite, we observed that genes with a steady increase in expression under cold deprivation conditions were mainly associated with cluster 3 and 6 (Fig. 9B). For the cluster 3, these paradormancy-associated genes are markedly up-regulated in prolonged cold deprivation while they slightly decrease in natural conditions (Fig. 4A). Surprisingly, a subgroup of genes activated during endodormancy (Cluster 6; Fig. 5A) was also up-regulated in the shallow dormancy stage (Fig. 9B). Overall, we found that most of the GO terms for the genes activated during shallow dormancy were associated with RNA metabolic processes, RNA splicing and regulation of gene expression (Fig. 4B, Fig. 5B). Furthermore, among the 116 marker genes previously identified for shallow dormancy (Fig. 8), we found several *LRR* kinases and *PavDAM1* (Supplementary Table 6) that were previously identified to be expressed during paradormancy and dormancy onset (Fig. 4C), but also candidate genes related to RNA modification such as a splicing factor (PAV06_REGINAg0287151). These results suggest a critical reprogramming of transcriptional processes under prolonged cold deprivation conditions.

## DISCUSSION

### Expected phenology disorders in fruit trees due to warmer temperatures in the context of climate change

Our experiments under constraint temperatures confirmed that sweet cherry reproductive cycle and phenological phases such as dormancy and flowering are strongly driven by temperature and we highlighted how dormancy progression could be impacted by none or low chilling accumulation. Sweet cherry tree appeared to be very sensitive to milder winter temperatures. Therefore a substantial chilling accumulation is crucial although a slight decrease in accumulated chill can be tolerated as observed for the trees submitted to a short cold deprivation, from July to November, which were able to progress through dormancy similarly to the control trees. We demonstrated that cold deprivation provokes phenological shifts such as delays in senescence and flowering but also leads to bud necrosis, bud default, lower quality and quantity of flowers. These results are reinforcing previous observations done on fruit trees which have reported phenological changes due to increasing temperatures (Atkinson *et al*., 2013). The delaying effects could be explained by warm temperatures during early dormancy phases just after bud set, as it has been described in beech and birch where similar delays in budburst were observed after warming treatments (Malyshev, 2020; Beil *et al*., 2021). Previous studies have also determined that sweet cherry trees have the capacity to postpone dormancy onset and maintain an active but reduced growth under short days at mild temperatures (Heide, 2008), in agreement with the shift in senescence we observed under cold deprivation. Moreover, flowering disorders, such as flower abortions, were reported on Japanese pear trees following a heat treatment in a greenhouse (Tominaga *et al*., 2022) while in apple, insufficient chilling led to physiological symptoms such as delayed foliation, low yield, and fruit physiological disorders (Rana et al, 2009; Legave *et al*., 2015).

Cold deprivation treatments gave a first glimpse of damages that could occur in the future due to how global warming will induce a decrease in cold accumulation during winter for sweet cherry trees. These experiments are also useful to better anticipate how temperatures will impact fruit trees and to assist breeders for the development of cultivars better adapted to future climatic conditions. For instance, our study under constrained temperature could allow defining safe winter chill values for projection of production areas under future climatic conditions (Campoy *et al*., 2019). We confirmed that, even under short day photoperiod, regular endodormancy is not induced at mild temperatures (21°C) in sweet cherry trees which may require low temperatures below 9°C for growth cessation (Heide, 2008).

To compensate the negative effect of climate, it has been proposed to cultivate low chill cultivars in combination with chemical agents used to promote bud break in moderately warm areas or to cultivate moderate or high chill cultivars in northern latitudes that are predicted to have an increase in the length of the growing season compatible with the presence of temperate fruit trees such as sweet cherry (Campoy *et al*., 2011). Nevertheless, in lower latitudes and warmer areas, only cultivation in mountainous areas could be appropriate to accumulate enough cold during winter period to ensure optimal flowering and fruit production.

### Molecular actors and signaling pathways involved in the regulation of dormancy in response to temperature

Many biological pathways have been previously reported to play a role in fruit tree dormancy progression such as metabolism activity, phytohormones, oxidative stress, transport capacity and transcription factors (Beauvieux *et al*., 2018). Our study provides new and reliable knowledge on these signaling pathways and genes implicated in dormancy progression by describing in details how constrained temperature can regulate their expression and could influence dormancy progression.

First, we suggest that genes highly expressed during paradormancy, i.e. before the seasonal temperature decrease, and subsequently down-regulated upon a cold signal, could be identified as dormancy onset regulators (clusters 3 and 5). They are involved in RNA metabolic and secondary metabolic processes, gene expression, cell communication and signaling, probably associated with high cellular activity and flower organogenesis to prepare the flower bud before winter dormancy, as previously described (Vimont *et al*., 2019). Under cold deprivation, some of these genes are steadily up-regulated, thus potentially maintaining cellular activity and preventing the induction of endodormancy. This is the case for example for key dormancy genes *PavDAM1* and *PavDAM6*, which are up-regulated or maintained at a high expression under cold deprivation conditions, respectively while they are promptly down-regulated by cold temperatures, with a timing correlated with senescence. These results may suggest that a cold-induced inhibition of *PavDAM1* and *PavDAM6* is necessary for the induction of endodormancy and related but not critical for growth cessation and senescence, therefore confirming their major role in paradormancy and dormancy onset (Li *et al*., 2009; Falavigna *et al*., 2019; Vimont *et al*., 2019). Interestingly, transgenic studies demonstrated that overexpression of Japanese apricot *PmDAM*6 in apple induced growth cessation and bud formation (Yamane *et al*., 2019), which suggest a complex regulation of dormancy onset by *DAM* genes.

While the expression of a majority of genes is inhibited after dormancy onset, marking a potential arrest of major signaling pathways, multiple studies including this one have revealed that specific genes are activated during endodormancy (Liu *et al*., 2012; Khalil-Ur-Rehman *et al*., 2017; Falavigna *et al*., 2019; Vimont *et al*., 2019; Canton *et al*., 2021; Prudencio *et al*., 2021; Calle *et al*., 2022). Their expression is potentially essential for buds to go through endodormancy so they can be characterized as dormancy maintenance regulators. Here, based on the response to constraints temperature conditions, we showed that these genes are induced upon exposure to cold temperatures and subsequently repressed at the end of endodormancy. This is the case for example for *PavDAM4*, which has been previously described as a key regulator for dormancy onset and maintenance (Zhu *et al*., 2020; Yu *et al*., 2021; Miyawaki-Kuwakado *et al*., 2024). In particular, Miyawaki-Kuwakado and colleagues (2024) estimated that an exposition to prolonged cold conditions is needed to suppress *DAM4* expression in Yoshino-cherry trees, in line with our observations that a certain amount of cold accumulation is necessary to induce then down-regulate the expression of endodormancy genes. Therefore, we may associate *PavDAM4* low expression levels under cold deprivation conditions to the non-induction of endodormancy, similarly to results obtained in Japanese pear where the repression of a *DAM* gene named *PpMADS13-3* by cold deprivation were associated to abnormal dormancy progression and flowering disorders (Tominaga *et al*., 2022).

Previous studies have reported a significant up-regulation of genes involved cellular activity, gametophyte development, photosynthesis and metabolic processes upon the endodormancy to ecodormancy transition (e.g. Liu *et al*., 2012; Vimont *et al*., 2019; Prudencio *et al*., 2021; Yu *et al*., 2021; Calle *et al*., 2022). In this work, we also identified clusters of genes displaying a high expression level after a prolonged cold signal exposure. These genes were mostly associated with growth resumption through cell cycle metabolic activities, and glycolytic processes, similarly to genes involved in cell cycle and mitotic cell cycle processes previously reported during the endodormancy to ecodormancy transition in peach and apricot (Yu *et al*., 2020). We found that they were maintained at low expression levels under cold deprivation conditions, associated with delayed dormancy release and bud break thus confirming their major role during the transition toward ecodormancy. This is the case for example for *PavCYCD3*, from the cell cycle-associated cyclin gene family (Menges *et al*., 2006), which was also found to be up-regulated at ecodormancy stage in Japanese pear (Saito *et al*., 2015). On another note, the resumption of glycolytic activity following dormancy release has been well-documented in several plant species and is closely associated with the mobilization of carbohydrate reserves. More specifically, the degradation of starch into soluble sugars such as glucose and fructose provides essential substrates for energy production and the production of amino acids, both of which are critical for flowering (Marafon *et al*., 2011; Stein and Granot, 2018; Chmielewski and Götz, 2022). In Japanese pear, insufficient exposure to chilling temperatures has been reported to disturb carbohydrate metabolism, potentially leading to developmental anomalies such as flower bud abortion or necrosis (Honjo, 2002; Marafon *et al*., 2011). These observations suggest that, following the satisfaction of chilling requirements, subsequent exposure to warmer temperatures during the ecodormancy phase may play a regulatory role by modulating the expression of genes involved in metabolic and cell activity pathways.

### Shallow dormancy as a specific molecular state induced by warm autumn and winter temperatures

The different levels of cold deprivation have been shown to gradually reduce deep dormancy to the point where we observed a particular dormancy stage characterized by a specific molecular state that we identified as shallow dormancy. It is different from the concept of low dormancy depth that have been especially proposed in forest trees (Søgaard *et al*., 2009; Malyshev, 2020; Garrigues *et al*., 2023). Dormancy depth is defined by the amount of heat accumulation needed to reach budburst throughout dormancy (Vitasse *et al*., 2014). It is species and cultivar-dependent and reaches a peak between October and December. In this context, low dormancy depth is defined when a tree is close to budburst following chilling accumulation, which may correspond to ecodormancy. In contrast, shallow dormancy is characterized by the inability to enter endodormancy in the absence of a cold signal, associated with incomplete dormancy progression and developmental anomalies.

The clustering analysis based on molecular data suggests that this shallow dormancy stage is associated but distinct from paradormancy. In the absence of cold signal to end paradormancy and induce endodormancy, we expected to observe maintenance of paradormancy. However, trees under cold deprivation entered a form of dormancy visibly marked by senescence and growth cessation, thus suggesting a physiological switch. Since previous studies have suggested that the timing of autumn senescence is correlated with spring phenology timing (Keenan and Richardson, 2015), we can hypothesize that endogenous signals always induce the initiation of growth cessation and a stage of minimal dormancy depth to ensure survival even through a very warm winter. This hypothesis is supported by the expression profiles of genes within the paradormancy Cluster 5 and ecodormancy Cluster 1, which exhibit a progressive downregulation beginning in summer, under all conditions from cold exposure to cold deprivation. These patterns suggest that signaling pathways are inhibited regardless of the temperature conditions, potentially under the control of internal inhibitors (Thomas and Vince-Prue, 1996; Battey, 2000). Overall, we found that downregulated genes were involved in metabolic processes, cell cycle, cytoskeleton organization, fatty and organic acid biosynthesis, thus confirming that cell activity and development are inhibited by endogenous signals to prevent growth under endodormancy or shallow dormancy equally.

On the other hand, results highlighted that shallow dormancy was a stage specifically characterized by the increased expression of genes involved in transcriptional regulation through RNA modification and post translational modification. This is in agreement with reports that environmental stresses can trigger rapid changes in gene expression by modifying RNA metabolism. Alternative splicing and in particular the regulation of pre-mRNA splicing have been reported to be induced by heat stress (Rosenkranz *et al*., 2022) and can play a role in stress acclimation of plants. For instance, RNA splicing, processing, transport and decay plays a major role in the molecular response to abiotic stress including temperature (Rosenkranz *et al*., 2022; Yan *et al*., 2022; Gururani, 2023; Zhang *et al*., 2023; Cai *et al*., 2025). The regulation of alternative splicing under elevated temperature undoubtedly plays a significant role in the stress acclimation of plants (Rosenkranz *et al*., 2022). Therefore, the gradual increase in RNA metabolism and transcriptional regulation could be one of the tree’s responses to control the balance between growth and survival during a prolonged cold deprivation, which can be considered as a major abiotic stress. However, whether RNA modification is beneficial for stress acclimation or detrimental due to the reduced capacity of the cell to synthesize proteins, in general (Rosenkranz *et al*., 2022) or for bud dormancy in particular, is not clear. Further investigation will be necessary to identify the targeted pre-mRNAs and understand the role the transcriptional regulation under prolonged cold deprivation.

## Conclusion

Our temperature-constrained experiments identified key genes involved in major signaling pathways—such as phytohormone signaling, oxidative stress response, transport, cell cycle, transcriptional regulation and carbohydrate and secondary metabolism—as central to the temperature-mediated control of dormancy in sweet cherry. Moreover, extreme cold deprivation induced a distinct shallow dormancy, providing insight into how warmer winters may affect phenological events. This shallow dormancy stage, defined by a unique molecular signature, may reflect an acclimation response to severe abiotic stress. While these findings offer a robust framework for understanding how dormancy progression is regulated by temperatures, functional validation in model or closely related fruit tree species will be necessary to confirm the key roles of these candidate genes for adaptation to future warmer winters.

## Supplementary data

Figure S1. Phenological changes induced by cold deprivation and exposure.

Table S1: Detail information on sequencing and mapping processes for each sample.

Table S2: Differentially expressed genes (DEGs) among 48, 133 genes identified in reference genome.

Table S3: GO enrichment analysis indicating Cellular Component (CC), Molecular Functions (MF) and Biological Processes (BP) over-represented in DEGs for each cluster compared to the entire gene set of DEGs.

Table S4. Predictive transcriptomic statistics.

Table S5: Gene list predicting shallow dormancy stage.

Table S6: DEGs expression under cold deprivation from August to April.

## Acknowledgements

We thank Xavier Lafon for managing the sweet cherry potted trees. The RNA sequencing was performed at GET-PLAGE sequencing facility in Campus INRAE of CASTANET-TOLOSAN. We are grateful to the Genotoul bioinformatics platform Toulouse Occitanie (Bioinfo Genotoul, https://doi.org/10.15454/1.5572369328961167E12) for providing computing resources. We warmly thank Anthony Bernard for his reviewing work.

## Author contributions

BW and MF designed the original research. MF performed the sampling and RNA-seq, and analyzed the RNA-seq data with BW. MF and HB monitored the trees’ phenology. SP developed the predictive transcriptomics model and script. MF and BW wrote the article with the assistance of all the authors. All authors have read and approved the manuscript.

## Data availability

RNA-seq data that support the findings of this study have been deposited in the NCBI Gene Expression Omnibus under the accession code GSEXXXX (data submitted in June 2025).

## Abbreviations

IPCC: Intergovernmental panel on climate change
DAM: Dormancy associated MADS-box
GA: Gibberellins
ABA: Abscisic Acid
ROS: Reactive oxygen species
RNA: Ribonucleic Acid
BBCH: Biologische Bundesanstalt Bundessortenamt und CHemische Industrie
CP: Chill portions
PVP-40: Polyvinylpyrrolidone average mol weight 40000
DNA: Deoxyribnucleic Acid
QC: Quality control
TPM: Transcripts per million
DEG: Differentially expressed genes
PCA: Principal component analysis
GO: Gene ontology
CHS: Chalcone synthase
LRR: Leucine rich repeat
GPX6: Gluthathione peroxidase X6
NCED9: Nine-cis-epoxycarotenoid dioxygenase 9
CYCD3: Cyclin D3
TUB8: Tubulin 8
PFK5: Phosphofructukinase 5
PAV: Prunus avium
MADS: MCM1,AG,DEFA,SRF
MF: Molecular functions
BP: Biological processes

